# Mating and starvation modulate feeding and host-seeking responses in female bed bugs, *Cimex lectularius*

**DOI:** 10.1101/2020.10.30.362806

**Authors:** Ahmed M. Saveer, Zachary C. DeVries, Richard Santangelo, Coby Schal

## Abstract

Adaptive insect behavior is subject to modulation by internal physiological states and external social contexts to enhance reproductive fitness and survival. The common bed bug, *Cimex lectularius* is an obligate hematophagous ectoparasite that requires host blood for growth, development, and reproduction. We investigated how mating, starvation and social factors such as harassment by males affect host-seeking, blood feeding, oviposition, and survival of female bed bugs. The percentage of females that fed and the amount of blood they ingested were greater in mated females (90–100%) than in unmated females (48–60%). Mating state also modulated the female’s orientation towards human skin odor in an olfactometer; more mated (69%) than unmated (23%) females responded to human odors. The response rate of unmated females (60%) to human odor increased with longer starvation period, while the opposite pattern was observed in mated females (20%). Although fecundity after a single blood meal was unaffected by long or short residence with males, females subjected to frequent copulations had lower survivorship and lifespan than females subjected to males for only 24 h. Taken together, these results indicate that behaviors are adaptively expressed based on the internal physiological state to maximize survival and reproductive fitness.

## INTRODUCTION

Nutritional and mating states can profoundly change the behavior of animals, including insects. Therefore, adult insects modulate their behavior in response to their metabolic needs and thus enhance reproductive fitness and survival. Previous studies have highlighted the modulation of behaviors by the mating and nutritional states in insect species with different ecology and life histories, including the fruit fly *Drosophila melanogaster* ^1,2^, Mediterranean fly *Ceratitis capitata* ^3^, lepidopteran species such as *Spodoptera littoralis* ^4^, and hematophagous insects such as *Aedes aegypti* ^5^ and the kissing bug *Rhodnius prolixus* ^6^. The emerging model is that mating and feeding events are tightly coordinated and drive the expression of specific behaviors, depending on the immediate needs of the insect. Understanding how these specific adaptive behaviors are controlled by the internal physiological state is critical to design new nonlethal strategies to target major insect pests ^5,7^.

The common bed bug, *Cimex lectularius* L. (Hemiptera: Cimicidae) is an obligate blood-feeding human ectoparasite that remains a public health pest worldwide ^8,9^. Bed bugs require frequent blood meals to support development, reproduction, and survival ^10^. Adult bed bugs are long-lived, and females require multiple feedings and matings to support multiple egg-laying cycles ^10,11^. The expression of these behaviors, to a large extent, depends on external conditions such as temperature, humidity, and host availability, and internal states including metabolic state, mating, starvation, and oocyte maturation ^10^. However, some behaviors can produce conflicting outcomes. While mating is required for oocyte maturation and oviposition of fertile eggs, social factors such as multiple matings, and harassment by males can be risky because extra-genital hemocoelic (traumatic) insemination in bed bugs can cause injury and transmit disease ^12,13^. Likewise, while blood-feeding is essential, feeding on host blood is risky, due to host defensive behavior ^14^ and increased post-feeding body volume of females attracts more males and thus higher mating rate, causing reduced female longevity ^15,16^. Therefore, it is fair to surmise that bed bugs should tightly regulate host-seeking, mating, and blood-feeding so these behaviors are coordinately expressed to maximize fitness while minimizing risk. State-dependent behavioral modulation has been documented in other hematophagous insects, including mosquitoes ^17^ and triatomines ^6,14^, but not in bed bugs.

When searching for a blood meal, bed bugs use host-associated cues, including heat ^18^, carbon dioxide ^19^, and odors ^20^. The integration of various host cues is poorly understood, but in an olfactometer bed bugs strongly respond to and orient towards human odors alone, independent of all other host-associated cues ^21^. Although many aspects of bed bug behavior related to host-seeking, blood-feeding, and mating have been documented ^12^, there is paucity in our understanding of how these behaviors are regulated by internal physiological states. For example, we recently showed that whereas mated females responded robustly to human odors 7– 10 days after ingesting a blood meal, unmated females did not respond in these bioassays ^21^.

We hypothesized that host-seeking and blood-feeding may depend on the females’ mating status. Moreover, we suspected that unmated females may process the blood meal more slowly than mated females and thus might be stimulated to host-seek after longer starvation periods. Our experimental design compared the percentages of females that blood-fed, size of the blood meal, and the human odor-mediated responses of unmated and mated female *C. lectularius* subjected to four starvation periods. We used a heated artificial feeder and a glass Y-tube olfactometer to assess feeding and orientation to host cues, respectively. In addition, we also measured fecundity and survival of females exposed to short- or long-term associations with males; the latter exposes females to harassment and potentially risky frequent copulations. Our results show that starved mated females are more stimulated to blood-feed and they ingest larger blood meals than starved unmated females. Moreover, starved mated females are strongly attracted to human odors, but attraction is dramatically reduced with longer starvation. We observed the opposite trend in starved unmated females, whose propensity to orient to human odors and take a blood-meal increased with longer starvation.

## RESULTS

### Starved unmated females ingest smaller blood meals than mated females

To explore the relationship between mating status and nutritional state, we fed virgin *C. lectularius* females, retained only fully engorged females, set them up into treatment groups of virgin (hereafter ‘unmated’) and mated females, and starved them for 8, 20, 30, and 40 days. For each experimental group, we examined the percentage that blood-fed and the size of their blood meals using an artificial feeder system. Whereas a high percentage of mated females fed (90– 100%), independent of the length of starvation and the size of their blood meals, significantly fewer unmated females blood-fed (48–60%) after 8 and 20-days of starvation (Fisher’s exact test (two-tailed): *P*<0.0001 and *P*=0.0001, respectively; Fig. 1a). The percentage of unmated females that blood-fed increased over time (75–97%) so that after 30 and 40-days of starvation there were no significant differences between mated and unmated females (Fisher’s exact test (two-tailed): *P*=0.112 and *P*=0.6139, respectively; Fig. 1a).

**Figure 1.**
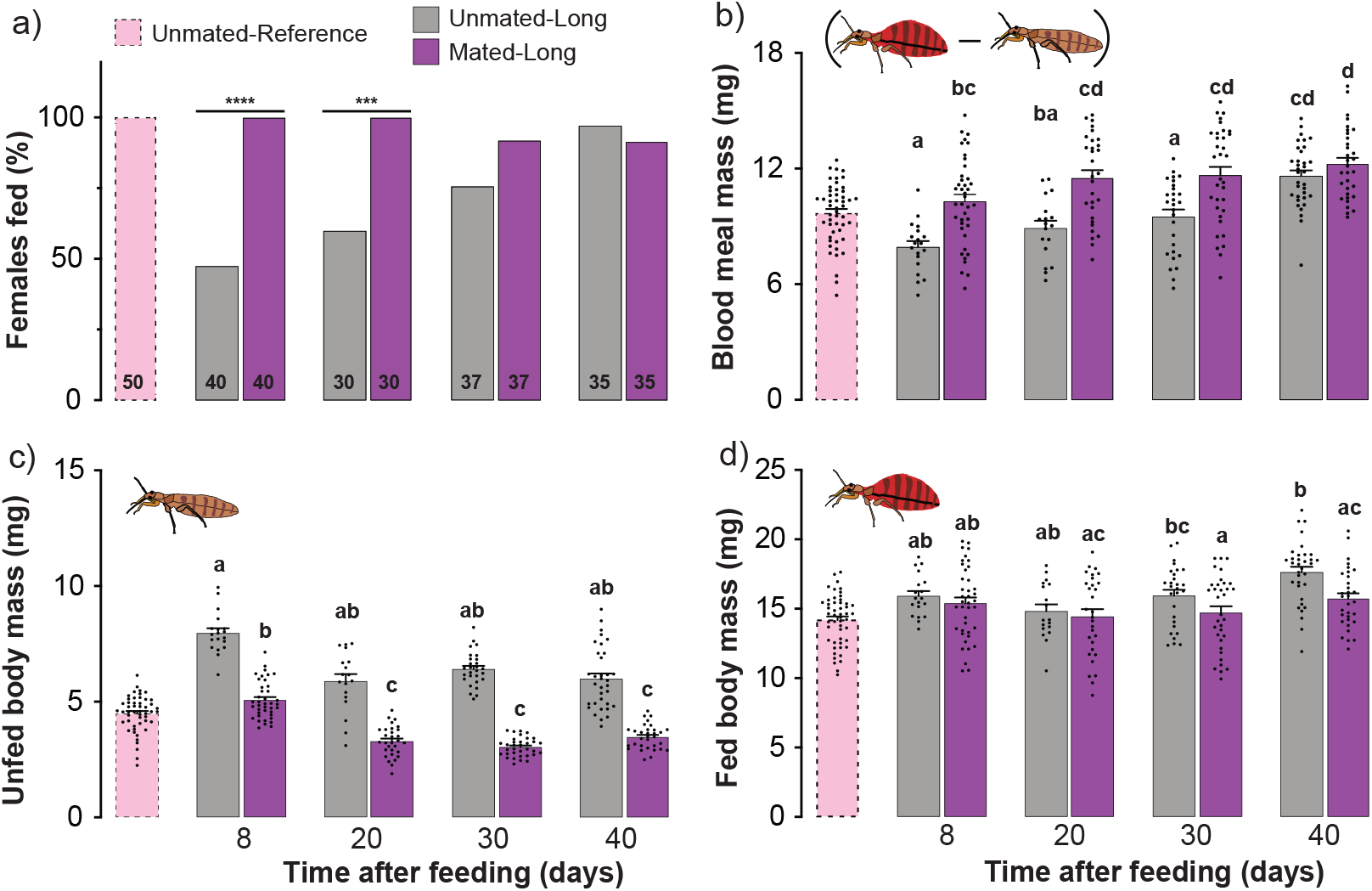
Effects of mating and starvation on the feeding parameters of *Cimex lectularius* females. (a) Percent feeding represents the percentage of tested females that ingested blood, determined visually by presence of blood in the digestive system. Unmated and mated female body mass before (b) and after (c) blood-feeding. Blood meal mass (d) was derived for each bed bug by subtracting its unfed body mass before feeding from its body mass after feeding. Females were allowed to fully engorge in a single blood meal within 8–10 days of eclosion and divided into three treatments. Unmated-Reference: not exposed to males; Unmated-Long: each female was housed with a sterile male (intromittent organ surgically ablated, denoted by an X within the male sign) until she died; and Mated-Long: each female was housed with a fertile male (denoted by a male sign) until she died. Both Unmated-Long and Mated-Long females were exposed to four starvation periods: 8, 20, 30 or 40 days. Numbers within bars represent the total number of replications for that treatment. The percent feeding differences between unmated and mated females for 8, and 20–days starvation periods are significant at **** *P*<0.0001 and *** *P*=0.0001, according Fisher’s exact test (two-tailed). Bars with different lower-case letters indicate significant difference (*P*<0.05) according to Kruskal–Wallis, Dunn’s post hoc test.

Likewise, 8, 20, and 30-days starved unmated females ingested significantly smaller blood meals than mated females (mean±SEM: 7.956±0.301, 8.919±0.381, and 9.535±0.363 mg, respectively; Kruskal-Wallis (Dunn’s multiple comparison) test: *P*<0.05; Fig. 1b), but their blood intake increased with longer starvation, with no difference between the two groups after 40 days of starvation. This difference is because unmated females retained most of the blood from their first blood meal, as indicated by their significantly greater unfed body mass than in mated females in all four starvation periods (Kruskal-Wallis (Dunn’s multiple comparison) test: *P*≤0.0001; Fig. 1c). In mated females that were starved for 8 days, the blood meal mass was only 2-fold their unfed body mass of 5.07 mg (mean±SEM: 2.065±0.077 fold; Figs. 1b, 2), likely because these females retained some blood from their previous blood meal and were still maturing oocytes (oviposition lasts for ~10 days after each blood-feeding). However, after 20 days of starvation the blood meal mass of mated females increased to 3.4-fold their unfed body mass of 3.28 mg (mean±SEM: 3.462±0.1546 fold; Figs. 1b, 2), as these females digested all their previously ingested blood and oviposited all their eggs. Blood meal mass continued to increase in mated females through 40 days of starvation (Fig. 1b).

In contrast, the blood meal mass of unmated females that were starved for 8 days was significantly smaller than in mated females (mean±SEM: 7.956±0.3017 mg; Kruskal-Wallis (Dunn’s multiple comparison) test: *P*=0.01; Fig. 1b), likely because of a larger unfed body mass of 7.96 ± 0.2033 mg (SEM), related to their slow digestion of the previous blood meal and retention of eggs (Fig. 1c). In these females, a smaller blood meal was sufficient to attain a large fed body mass of 15.92 ± 0.3452 mg (Fig. 1d). The ratio of blood meal mass to unfed body mass increased to only 2-fold and was not significantly different among the 20, 30, and 40-day starvation groups (Kruskal-Wallis (Dunn’s multiple comparison) test: *P*>0.99; Fig. 2). This pattern represented larger blood meals and increasingly smaller body mass as the previous blood meal was slowly digested (Fig. 2).

**Figure 2.**
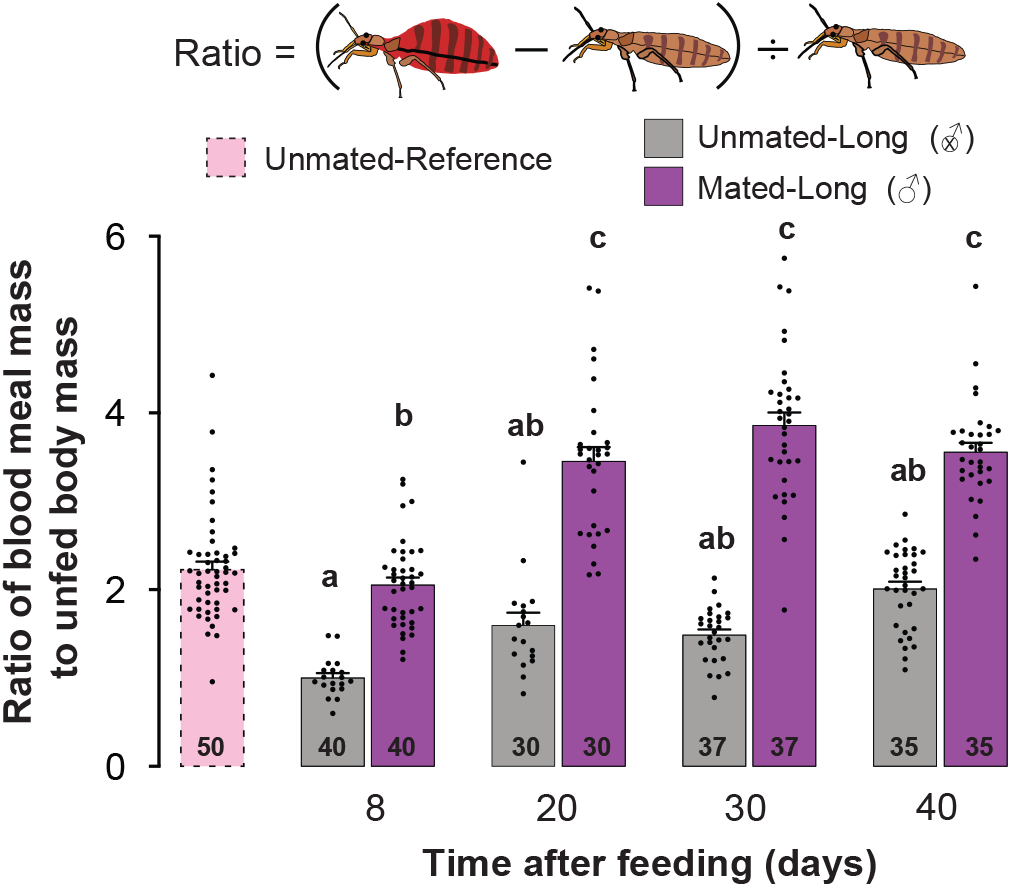
Effects of mating and starvation on the size of the blood meal of *Cimex lectularius* females. The ratio of blood meal mass to the unfed body mass of three treatment groups. Unmated-Reference (not exposed to males); Unmated-Long: each female was housed with a sterile male (denoted by an X within the male sign) until she died; and Mated-Long: each female was housed with a fertile male (denoted by a male sign) until she died. Both Unmated-Long and Mated-Long females were exposed to four starvation periods: 8, 20, 30 or 40 days. Numbers within bars represent the total number of replications for that treatment. Bars show mean±SEM (*n*=30–50) and all replicates are displayed. Bars with different lower-case letters indicate significant difference (*P*<0.05) according to Kruskal–Wallis, Dunn’s post hoc test.

For reference, we also assessed the same parameters in a group of virgin females that took their first adult blood meal within 8–10 days after eclosion, and were never exposed to males after the eclosion. These females behaved more like mated starved females than unmated starved females. The percentage of females that fed (Fig. 1a), their blood meal mass (Fig. 1a, b, c), and the ratio of blood meal mass to unfed body mass (Fig. 2) were similar to the respective parameters of mated females, and much higher than for unmated 8-day starved females.

### Mating and starvation modulate orientation to host odors

To investigate the effects of mating and starvation on orientation to human odor, we deemed it important to control for recurrent copulations and the presence of males in the mated group. Therefore, we conducted olfactometer assays to determine the effects on starved females of being housed with fertile males (i.e., mated females) vs. sterile males (male with a surgically ablated intromittent organ) that could not copulate (i.e., unmated females); females in both groups experienced social interactions. When these conditions persisted throughout the female’s adult life (i.e., ‘Long’ treatments), significantly more mated than unmated females responded when starved for 8- or 20-days (Fisher’s exact test (two-tailed): *P*<0.0003 for both; Fig. 3a). Strikingly, responses of 40-day starved females were reversed, with significantly more unmated than mated females showing robust upwind orientation by walking towards the human skin swab (Fisher’s exact test (two-tailed): *P*<0.0086; Fig. 3a). Overall, the response rate of unmated females towards a skin swab increased with longer starvation, whereas the response rate of mated females declined with longer starvation.

**Figure 3.**
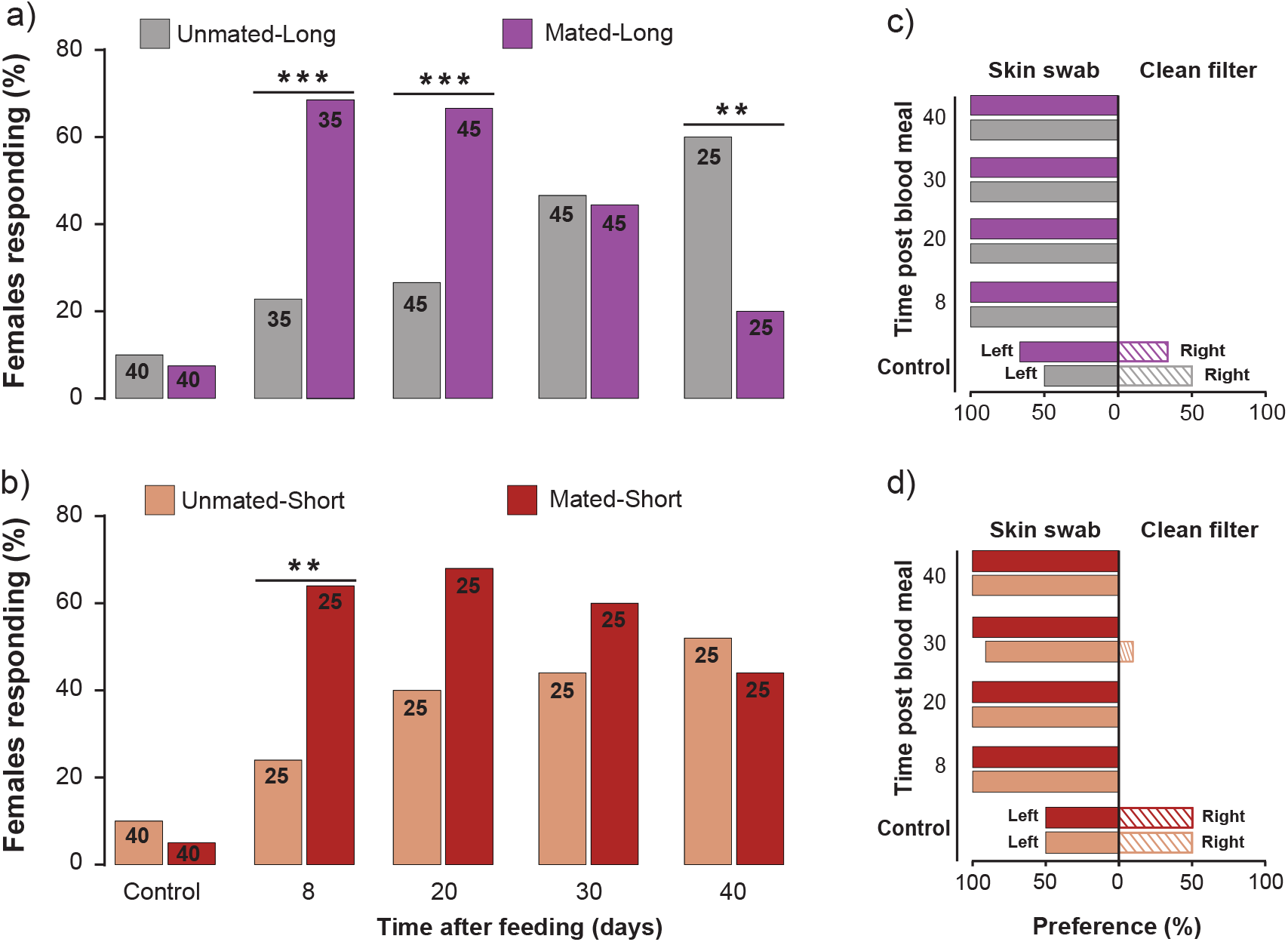
Effects of mating and starvation on orientation of female *Cimex lectularius* to human skin odor in an olfactometer. Percent response (a and b) represents the percentage of females that made a choice (skin swab or clean filter) of all females tested. (a) Unmated-Long: each female was housed with an infertile male (intromittent organ ablated, denoted by an X within the male sign) until she died; Mated-Long: each female was housed with a fertile male (denoted by a male sign) until she died. (b) Unmated-Short: each female was housed with an infertile male for 24 h and then with a female (denoted by an X within the male sign and then a female sign); Mated-Short: each female was housed with a fertile male for 24 h and then with a female (denoted by a male sign and then a female sign). (c) and (d) Olfactometer preference for human skin odor (left) vs. clean filter paper (right). Assays were conducted on four female treatment groups 8, 20, 30, and 40 days after they ingested a blood meal. Numbers within bars represent the total number of replications for that treatment. In (a) and (b), Control bioassays with clean filter papers of females of different starvation periods were pooled due to low response rates (<15%, *n*=40), and differences between unmated and mated females are significant at ** *P*<0.0096, *** *P*<0.0003, according to Fisher’s exact test.

A similar response pattern was apparent in females transiently exposed to males for only 24 h (i.e., ‘Short’ treatments). The orientation responses of unmated females gradually increased with longer starvation, whereas in mated females response rate increased and then declined; we found a significant difference between them only at 8 days of starvation (Fisher’s exact test (two-tailed): *P*<0.0096; Fig. 3b). However, we did not observe the sharp decrease in the response rate of mated females after 30- or 40-days of starvation that was apparent in the ‘Long’ experimental condition.

Females in all treatments were highly adept at detecting and preferably orienting toward human skin odor over the clean control filter paper, irrespective of their mating status, social conditions, nutritional state, and response rate in the olfactometer (Fig. 3c, d). Control assays with clean filter papers in both arms of the olfactometer resulted in low response rates (<15%) and random orientation to the left and right arms of the olfactometer. Of note is that although we defined a “response” and “making a choice” as reaching 1.5 cm into either arm of the olfactometer, 100% of the females that responded reached the respective filter paper.

### Female oviposition and survival

We hypothesized that long co-habitation with a fertile male would reduce female fecundity after a single blood-meal and lifetime survival because of constant harassment and physical damage to the female during repeated hemocoelic copulations. However, we found that the number of eggs oviposited during the first oviposition cycle (within 10 days after the blood-meal) was the same in both female groups (with a fertile male throughout vs. with a fertile male for only 24 h) and their cumulative mean numbers of eggs per female were similar (Fig. 4a, b). In both groups, females initiated egg laying on day-4 and ceased on day-10 post blood-feeding.

**Figure 4.**
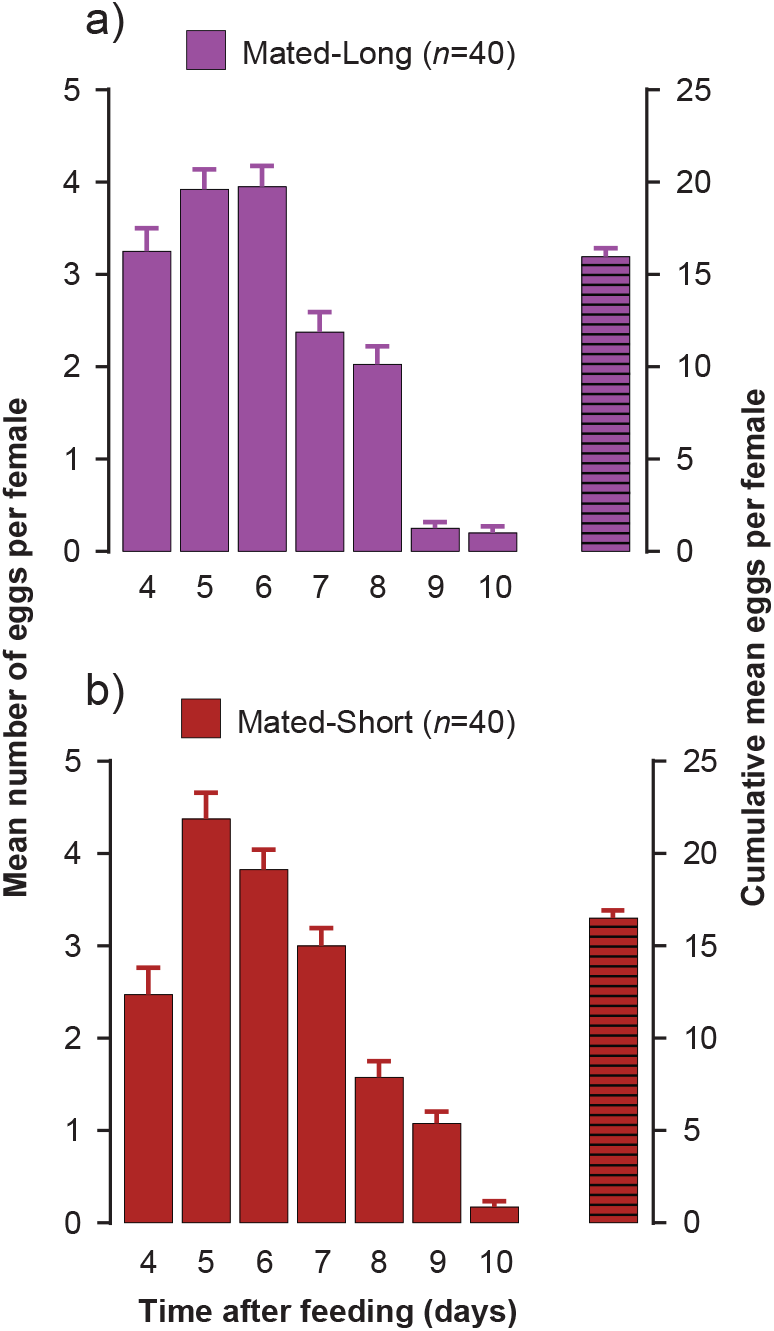
Egg production of female *Cimex lectularius* in the first oviposition cycle after a blood meal. (a) Mated-Long: each female was housed with a fertile male until she died (denoted by a male sign). (b) Mated-Short: each female was housed with a fertile male for only 24 h and the male was replaced with an antennaless female (denoted by a male sign and then a female sign). Females were monitored daily for the number of eggs they oviposited. Bars show mean daily eggs per female and cumulative mean number of eggs oviposited per female (±SEM, *n*=40).

On the other hand, survival was significantly affected, with mated-short females (each with a fertile male for only 24 h) surviving for a median of 55.5 days, whereas median survival of mated-long females (each exposed to a fertile male throughout her lifetime) was only 35 days (Log-rank (Montel-Cox) test: df=1; Hazard ratio (log-rank): 4.012; *P*<0.0001; Fig. 5). The same pattern was evident in unmated females: The median survival times of unmated-short (24 h with a sterile male) and unmated-long females (lifetime with a sterile male) were 111 days and 73 days, respectively (Log-rank (Montel-Cox) test: df=1; Hazard ratio (log-rank): 5.055; *P*<0.0001). Overall, mated females died at a significantly higher rate than unmated females (Log-rank (Montel-Cox) test: df=3; *P*<0.0001; Fig. 5), suggesting that both matedness and long-term presence of either a fertile or infertile male reduced female survival.

**Figure 5.**
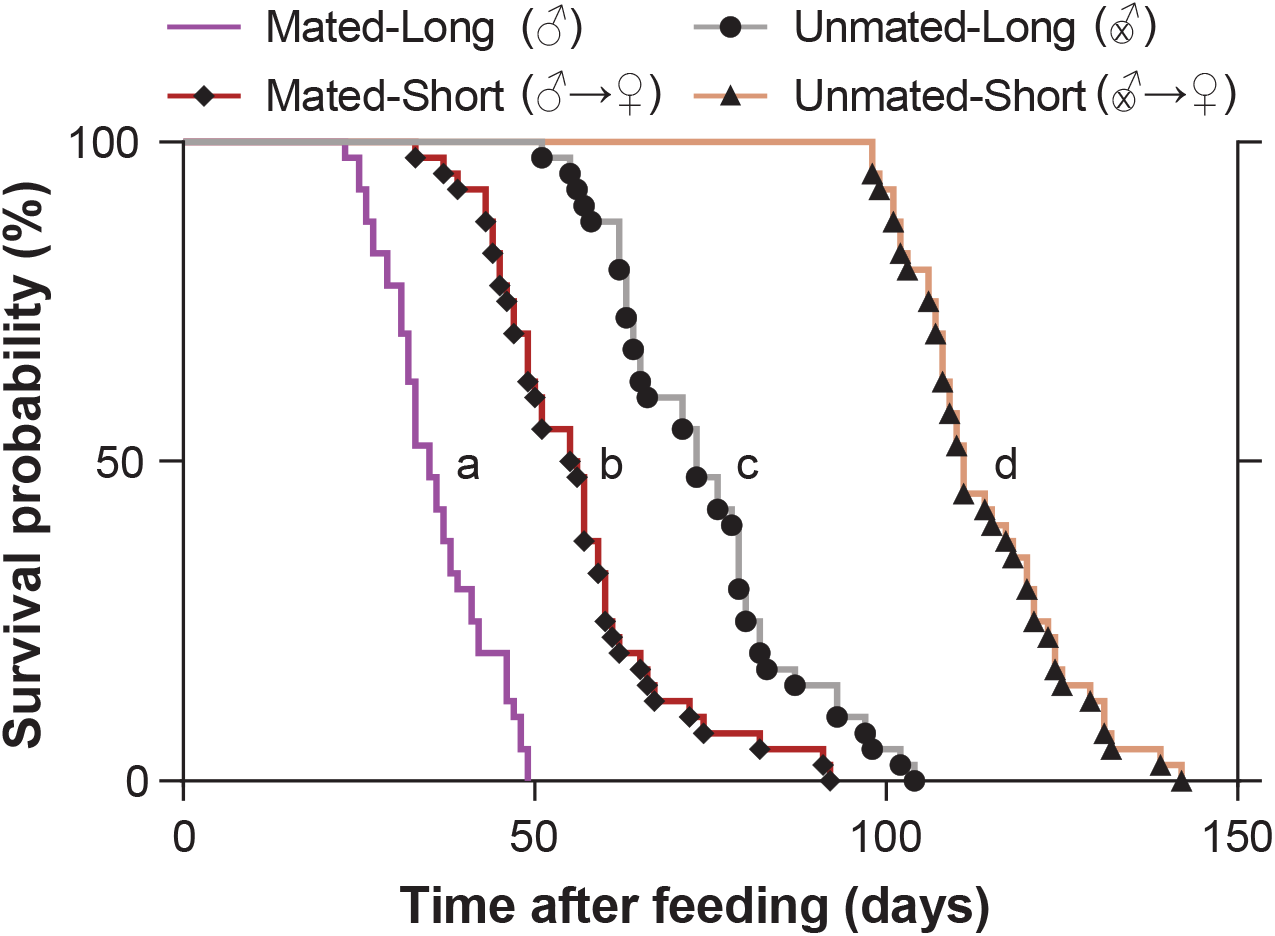
Effects of mating frequency and starvation on the survival of female *Cimex lectularius*. The four treatments include, Unmated-Long: each female was housed with a sterile male (intromittent organ surgically ablated, denoted by an X within the male sign) until she died; Mated-Long: each female was housed with a fertile male until she died (denoted by a male sign); Unmated-Short: each female was housed with a sterile male for 24 h and the male was replaced with an antennaless female (denoted by an X within the male sign and then a female sign); and Mated-Short: each female was housed with a fertile male for 24 h and the male was replaced with an antennaless female (denoted by a male sign and then a female sign). Survival curves with different lower-case letters are significantly different (*P*<0.0001) according to log-rank (Mantel-Cox) test.

Finally, we discovered a female “refusal” posture, whereby the female curled the ventral side of the abdomen, protecting the ectospermalege, a specialized cuticular region that guides the male phallomere, from males (Fig. 6a, supplementary video S1). This behavior was expressed only in the unmated-long and mated-long treatment females but not in the respective short treatment females (Fig. 6b), consistent with the idea that the “refusal” posture is expressed more in the presence of males.

**Figure 6.**
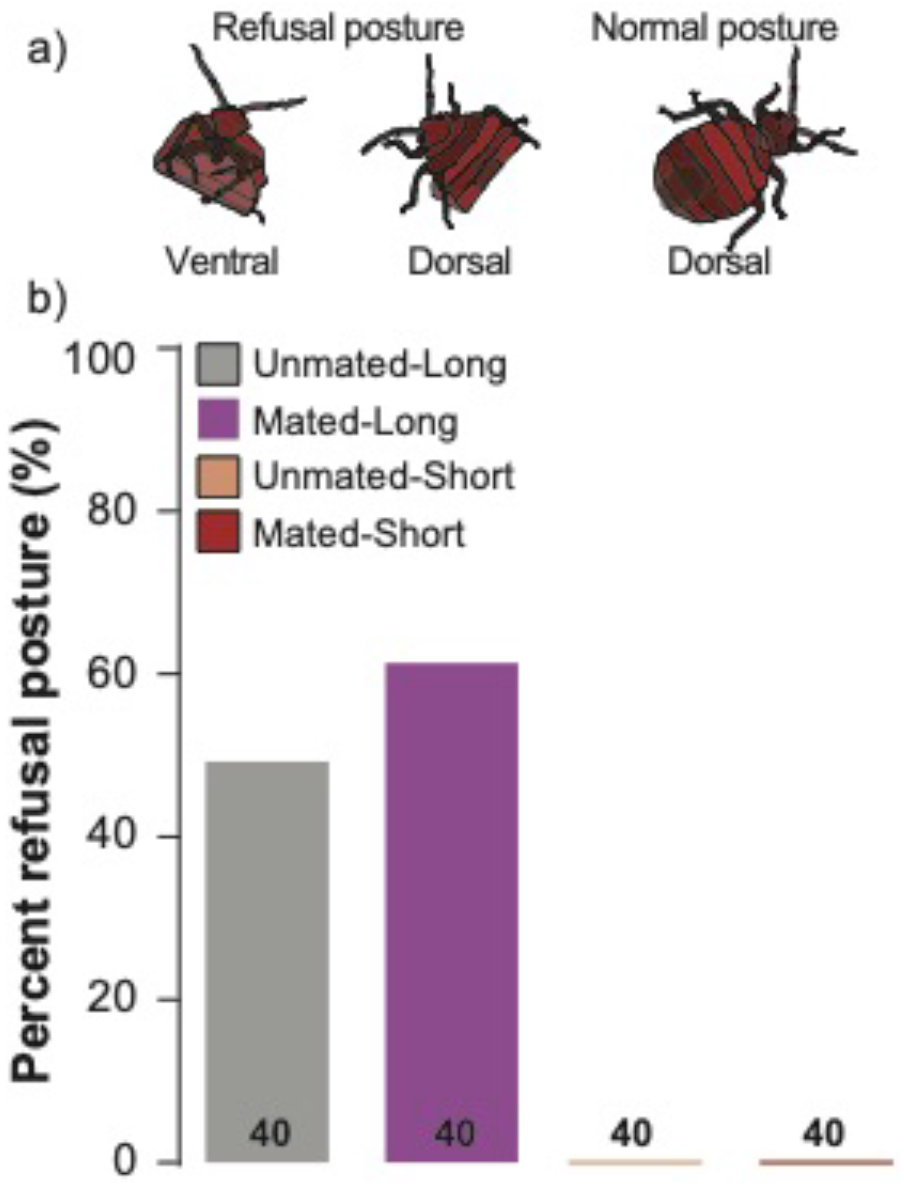
Expression of a female refusal behavior in *Cimex lectularius* under various mating and starvation conditions on the survival of female *Cimex lectularius*. (a) Dorsal and ventral views of the refusal behavior, and a female in a normal position. (b) Refusal behavior frequency in four treatments, showing the percentage of females in each treatment that expressed the refusal behavior at least once in their lifetime. Unmated-Long: each female was housed with a sterile male (intromittent organ surgically ablated) until she died; Mated-Long: each female was housed with a fertile male until she died; Unmated-Short: each female was housed with a sterile male for 24 h and the male was replaced with an antennaless female; and Mated-Short: each female was housed with a fertile male for 24 h and the male was replaced with an antennaless female. Numbers within bars represent the total number of replications for that treatment.

## DISCUSSION

The bed bug *Cimex lectularius* is an obligate ectoparasite, a lifestyle that requires the coordination of diverse behavioral repertoires, including host-seeking, blood-feeding, mating, oviposition, and aggregation to sustain development, reproduction, and survival ^10,12^. In support of the appropriate coordination of the sequence of these behaviors, adult female bed bugs are expected to monitor and integrate their changing needs and circumstances to express specific behaviors that increase reproductive fitness and minimize fitness costs. Indeed, while developing and validating a vertical Y-olfactometer for bed bugs, we observed a mating-dependent behavioral shift where 47% of mated females responded to human skin odor, but none of 20 unmated females responded to the same olfactory stimuli ^21^. In the present report, we followed-up on this observation because, to our knowledge, no experimental studies have investigated whether bed bugs modulate host-seeking and blood-feeding behaviors with changes in their physiological state such as mating and nutritional state.

Consistent with our previous results ^21^, we found that 64–69% of mated females responded to human skin odor 8 days after their last blood meal (Fig. 3). In contrast, only 24% of unmated females responded, revealing again that host-seeking behavior is profoundly modulated by the mating state of the bed bug. We observed higher host-seeking response rates in this study than in the previous report ^21^, possibly because we used slightly different durations of starvation and olfactometer conditions.

We speculated that the behavioral differences between unmated and mated females were related to the rate of processing of the blood meal, which would be affected in turn by the rate of oocyte maturation and oviposition. Specifically, we hypothesized that because unmated females retain their eggs, they have lower nutritional requirements and reduced metabolic rate ^22^ and therefore engage in less host-seeking and blood-feeding. To test this hypothesis, we extended the starvation period for both mated and unmated females. We found that the length of starvation had different effects on the host-seeking and feeding responses of mated and unmated females. Whereas host-seeking gradually increased in unmated females (24 to 60%) through 40 days of starvation, mated females became less responsive to host cues with longer starvation. These observations were consistent with the blood-feeding assays – with longer starvation, more unmated females fed and they took larger blood meals, even though they did not fully process some of their previous blood meal, as evidenced by their unfed body mass (Fig. 1b).

These results are consistent with our hypothesis that the differences between mated and unmated females are driven by the accelerated egg maturation and oviposition cycle of mated females. Mated females initiated egg laying 4 days after ingesting a blood meal, oviposited on average 15.3 eggs per female, and ceased oviposition 6 days later, 10 days after ingesting a blood meal (Fig. 4). Thus, 8 days after a blood meal, mated females digested most of the blood (Fig. 1b), oviposited most of their eggs, and became highly motivated to host-seek to support a second oviposition cycle. Indeed, both laboratory and field observations showed that, given the opportunity, mated females accept a second blood meal while the first blood meal is still being digested and females feed every 2.5 days on average ^10,12,23^. This is in contrast with other hematophagous insects, such as mosquitoes, where female host-seeking and feeding are suppressed for several days after a blood meal, until she completes laying a batch of eggs ^17,24^.

The strategy of unmated females was to feed little when the host is available, but with longer starvation periods, they became more stimulated to host-seek and ingest increasingly larger blood meals. This strategy is likely driven by the much-reduced nutritional demands of arrested egg maturation, which allow unmated females to digest the blood meal more slowly and use it for somatic maintenance rather than egg maturation. Metabolic differences between mated and unmated females support this hypothesis, with unmated females having reduced metabolic rates after feeding compared with mated females ^22,25^. Overall, this strategy would result in fewer host-seeking forays and less blood ingested by unmated females (Fig. 1b, c), until they mate and are stimulated to mature and oviposit fertile eggs. A similar strategy appears to operate in the closely related kissing bug *R. prolixus*, where virgin females remained unresponsive or even repelled by host-associated cues (CO_2_ and heat) until 20 days after ingesting a blood meal ^14^. *Rhodnius* is a much larger hemipteran than *C. lectularius*, it takes larger blood meals, and likely requires more time to digest the blood.

Surprisingly, host-seeking declined in mated females that were starved for 30 or 40 days, and this was especially apparent in females housed with fertile males (mated-long treatment) (Fig. 3). Two factors might account for this observation. The first is female aging and senescence, as 40 days of starvation in these females was beyond the 35-day median survival of females in this treatment group, and 100% of these females died by day 49 (Fig. 5). Thus, the females that we bioassayed 40 days after they ingested a blood meal were likely weak and less responsive to olfactory stimuli. This reduction in the host-seeking and blood-feeding responses of older mated females might be associated with their higher metabolic rate, senescence, or aging that could negatively affect olfactory responses, as shown in the *D. melanogaster* ^26^.

The second factor that likely underlies their early senescence is the unusual extra-genitalic, hemocoelic (traumatic) insemination in *C. lectularius*. Females housed with fertile males would receive multiple copulations that represent constant harassment, stress, and physical damage. These interactions with sexually aggressive males reduced their median adult lifespan by 63% (mated-long treatment) relative to females that were housed with fertile males for only 24 h and then with another female (mated-short treatment) (Fig. 5). These findings are consistent with previous studies in bed bugs on the adverse effects of multiple copulations on female survival ^12,13,16,27,28^. We also observed that mated-long females assumed a “refusal” posture, protecting the ectospermalege from the males (Fig. 6a, supplementary video S1). Nevertheless, males are able to circumvent these defensive postures by puncturing the intersegmental membrane away from the specialized ectospermalege, and thus cause substantial damage to the female. This injurious effect of males on females likely shapes female behavioral responses in the field that were constrained under our experimental conditions. For example, mated females might leave aggregations to avoid males, as demonstrated experimentally ^29^. Mated females might also seek refugia that are too narrow to accommodate themselves as well as courting males.

The lifespan of mated females was significantly reduced by both the long- and short-term presence of fertile males that represented high and low mating rates, respectively. Interestingly, however, we found no difference in female fecundity in the first oviposition cycle (~10 days post blood meal) between these two treatment groups (Fig. 4). Although our experiments were restricted to a single feeding and only one oviposition cycle, our results were similar to previous findings where higher mating rate dramatically reduced female lifespan with no significant effects on lifetime egg production ^27^. These findings suggest that mated females preferentially direct resources to egg maturation at the risk of significantly reduced lifespan, a strategic decision females make based on their physiological state and environmental conditions ^30^

Remarkably, we also detected a significant effect on females of non-copulatory harassment by males. Females housed with a male that could not copulate (intromittent organ ablated) for only 24 h and then with another female (unmated-short treatment) lived to a median age of 111 days (100% dead by day 142), whereas females housed continuously with an infertile male (unmated-long treatment) lived to a median age of only 73 days (100% died by day 104). This 34% decline in expected longevity, independent of copulation and egg production, can be attributed to male-specific harassment (Fig. 5). Males engage in a stereotyped behavior where the male repeatedly mounts the female’s dorsum, bends his abdomen to her ventral surface, and probes the female’s sternites with the paramere (intromittent organ). We observed that 50% of the unmated-long and 62.5% of mated-long females exhibited a ‘refusal’ posture at least once in their lifetime thereby making the ectospermalege inaccessible to males, while none of the unmated-short or mated-short females displayed this behavior (Fig. 6b). The expression of this refusal behavior in virgin females that obviously need to mate is particularly surprising, suggesting that male harassment is especially detrimental to the metabolic budget of starved females that feed less and endeavor to conserve energy. Unfortunately, our experimental design did not enable us to determine whether co-habitation with a female also would decrease survivorship of starved females relative to solitary females. It is possible that general disturbance of the starved female causes her to move and expend energy, which in turn reduces her lifespan. In this context, it is worth noting that by adding a female to the mated-short treatment, after the male was removed, the presence of the extra female might have decreased survivorship and thus resulted in underestimating the difference between the mated-long and mated-short treatments.

## Conclusions

Mating in female bed bugs strongly influences their host-seeking and blood-feeding behaviors, and the size of their ingested blood meal. Whereas mated females were highly stimulated by host odors to host-seek, and they ingested large and frequent blood meals, unmated females were less driven to seek a host and they ingested smaller and less frequent blood meals. The length of starvation also affected mated and unmated females differently. Host-seeking declined over starvation time in mated females, likely due to mating-induced physiological changes, whereas host-seeking responses in unmated females gradually increased. To our knowledge, this is the first report showing that multiple matings impair host-seeking in older starved female bed bugs, and the first to show that non-copulatory male harassment dramatically reduces female lifespan. Finally, our results show that female bed bugs that respond in the olfactometer orient towards human odor regardless of their mating and nutritional status.

## MATERIAL AND METHODS

### Insects

*Cimex lectularius* used for this study was the Harold Harlan (Harlan) population, originally collected in 1973 at Fort Dix, NJ, USA, and maintained in the laboratory. Colonies were maintained in an incubator at 27°C, approximately 30–50 % relative humidity (RH) and under a 12 h light:12 h dark cycle (lights off at 0800). Bed bugs were fed defibrinated rabbit blood (HemoStat Laboratories, Dixon, CA) using an artificial feeder system. A heated water bath (B. Braun Biotech Inc., Allentown, PA) circulated warm (34°C) water through a custom-made water-jacketed glass feeder. Blood in each glass feeder was retained with a stretched plant grafting tape (A.M. Leonard Horticultural Tool and Supply Co., Piqua, OH) on the bottom and ~4 mL of blood was introduced through the opening at the top. Bed bugs were housed in a polystyrene wide-mouth threaded colony container (Consolidated Plastics Company, Inc., Stow, OH) constructed by replacing the bottom with a nylon screen (0.3 mm mesh opening; BioQuip Products Inc., Rancho Dominguez, CA) and a piece of folded manila paper was provided so bed bugs could walk up to the screen and feed. Colonies were fed weekly by placing each container screen side up under a feeder for 15–20 min.

### Experimental design

To obtain unmated females, freshly fed 5^th^ instars were individually placed into glass vials (7.5 mL), each with a paper strip. Adult unmated females were collected on the day of eclosion and up to two days after eclosion and grouped in a polystyrene wide-mouth threaded container with a screened bottom, through which bed bugs could feed as described above. The females were group-fed to satiation within 8–10 days and each placed in a clean glass vial (7.5 mL) with a paper substrate. The following day (24 h after feeding) fully engorged individual females (9-11 days post eclosion) were subjected to one of the following four experimental treatments: 1) ‘Mated-Long’: each female was housed with and copulated frequently with a single normal male until she died; 2) ‘Unmated-Long’: each female was housed with a male with a surgically ablated intromittent organ (hereafter ‘sterile male’) that could not copulate, until she died; 3) ‘Mated-Short’: each female was housed and allowed to copulate with a fertile male for only 24 h and the male was then replaced with an antennaless female (to differentiate her from the experimental female); and 4) ‘Unmated-Short’: each female was housed for 24 h with a sterile male that could not copulate, and the male was replaced with an antennaless female. In the ‘Long’ experimental treatments, females experienced repeated harassment by males and either frequent successful copulations (mated-long) or unsuccessful mating attempts (unmated-long). In the ‘Short’ experimental treatments, females experienced lower harassment and either successful copulation during only 24 h with a male (mated-short), or unsuccessful mating attempts for 24 h (unmated-short). In the ‘Short’ treatments, males were replaced with antennaless females after 24 h to maintain social conditions that might affect reproduction and longevity ^31^. Males were 4–6 days post blood-feeding. We replaced dead males in two treatments; one in an unmated-long replicate and one in a mated-long replicate. Likewise, a dead antennaless female was replaced in one unmated-short and one mated-short replicate. Oviposition and female survival were monitored daily.

### Female ‘refusal’ behavior

We visually recorded female ‘refusal’ behavior as a result of starvation and male mating harassment at least once daily until the death of the female, in all four experimental treatments: Mated-Long, Unmated-Long, Mated-Short, and Unmated-Short. We scored ‘presence’ of refusal behavior if a female curled the ventral side of the abdomen, protecting the ectospermalege from males as illustrated in Fig. 6a and supplementary video S1.

### Blood-feeding assay

We assayed the effects of mating and starvation on the female’s propensity to blood-feed and the size of her blood meal using the unmated-long and mated-long treatments (*n*=30–40 per treatment). All the feeding experiments were performed between 3–6 h after onset of the scotophase (under red-light) and at 26 ± 1°C. At the beginning of the scotophase (1 h before testing), each female was weighed (MP8-1, Sartorius, Goettingen, Germany) and placed in a glass vial (7.5 mL) containing a folded paper strip leading to a nylon screened lid (0.3 mm mesh opening), so she could walk up to the screen and feed. Then, two artificial glass feeders (described above) were prepared side by side, and four vials were placed randomly under each feeder for 10 min – two vials were of unmated-long females and two mated-long females of one of four starvation periods (8, 20, 30, and 40 days). Immediately after the assay, females were cold-immobilized on an ice tray. At the end of the experiment, all the females were visually inspected for blood in their digestive system and females that did not feed were excluded from subsequent analyses that required a quantitative measure of blood-meal size. Females that ingested blood were weighed, independent of the size of their blood meal. Notably, the identity of each female was known throughout the experiment. We also prepared an additional group of unmated females, collected from a different cohort of 5^th^ instars, termed the ‘reference’ feeding group (*n*=50), representing the feeding propensity and the size of the first blood meal of virgin females within 8–10 days of eclosion. The reference group was prepared identically to the other groups except females were not exposed to males after eclosion. Females in these three treatment groups (mated-long, unmated-long, and reference) were not used in other experiments.

### Human skin swab collection

We used a filter paper to obtain human skin odors to test bed bug attraction to human odors, as previously described by DeVries et al. (2019). Skin swabs were collected in accordance with guidelines and regulations, and the experimental protocol was approved by a member of the ethics review committee under an approved North Carolina State University Review Board protocol (#14173). Informed consent for study participation was obtained from each human subject. The participant was requested to refrain from drinking alcohol, eating spicy foods, avoid the use of skin care products and any strenuous physical exercise, take a morning shower with a perfume-free shower gel (Cetaphil, Galderma Laboratories, L.P., Fort Worth, TX), and obtain the skin swabs 4–8 h after showering. Skin swabs were prepared with #1 Whatman filter papers (90 mm diameter, Whatman plc, Madistone, UK) as follows: 1) Rinse hands with tap water and dry before swabbing, 2) swab the left arm from hand to armpit for 12 sec and the left leg from the lower thigh to ankle for 12 sec using both sides of the same filter paper, 3) swab the left armpit for 6 sec using both sides of the filter paper, 4) place the filter paper into a 20 mL glass vial, and 5) repeat the steps on the right side of the body with a new filter paper. Skin swab samples were stored in a −30°C freezer. We used only one participant’s skin swabs for all the bed bug orientation experiments.

### Olfactometer bioassays

We used a vertical Y-tube custom-made glass olfactometer (14 mm inner diameter) to test the effect of mating and starvation on the orientation of female bed bugs to skin odors, as previously described ^21^. Charcoal filtered, humidified, medical-grade breathing air was pushed through the olfactometer at 300 mL min^−1^. A nylon screen walkway (Nitex, 300 μm, BioQuip Products Inc., Rancho Dominguez, CA) in the shape of the Y-tube was inserted in the olfactometer, serving as a walkway because bed bugs do not walk well on glass surfaces. Bed bugs were kept individually in 5 cm long ‘releasing tubes’ in the experimental room for 1 h before testing. Releasing tubes were made of centrifuge tubes, with the bottom removed and replaced with a nylon screen, so that air could flow through the tube but the bed bugs could not escape. Bed bugs were acclimatized by attaching the releasing tube to a separate air stream (300 mL min^−1^) for at least 5 min. After the acclimation period, 1/16th (4 cm^2^) of a skin swab filter paper was placed in a glass ‘odor pot’ connected with a ground glass fitting to the distal end of one arm of the olfactometer. We used 1/16th (4 cm^2^) of a skin swab filter paper as this size was sufficient to elicit robust odor-mediated responses in the male and female bed bugs ^21^. An identical odor pot with an identically sized clean filter paper (control) was connected to the opposite arm. A releasing tube with an acclimated female bed bug was attached to the downwind end of the olfactometer and observed for 5 min. The following behaviors were recorded: Bed bugs that moved upwind and made a choice by reaching 1.5 cm into either arm (odor or control) of the olfactometer were considered responders; bed bugs that failed to reach either of these points were considered non-responders. The positions of the skin swabs were randomized by switching the odor source to the right or left arm of the olfactometer. Control tests were performed with clean filter papers in both arms of the olfactometer to identify any positional bias. After each bioassay trial, the skin swab and clean filter paper were replaced. Each bed bug was used only once in the bioassay and all the experiments were performed between 3–6 h after onset of the scotophase under a single fluorescent light encased in a red photographic filter, illuminated from 1 m above the olfactometer. Olfactometer assays were performed on four female treatments – unmated-long, mated-long, unmated-short, and mated-short (*n*=25–45 per treatment) – of the following four starvation periods: 8, 20, 30, and 40 days. Bioassays with control filter papers in both sides of the olfactometer were conducted simultaneously with each treatment (*n*=10 per treatment). The glass olfactometers were cleaned with acetone and the nylon screen ramps were replaced after every 1–2 bioassays.

### Data Analysis

Fisher’s exact test (two-tailed) was performed to compare the effects of mating and starvation on the feeding propensity of females between treatments of the same starvation period. ‘Blood meal mass’ was calculated as (Fed body mass – Unfed body mass). The ratio of blood meal mass to unfed body mass was calculated as (Fed body mass – Unfed body mass)/Unfed body mass. The non-parametric Kruskal–Wallis test followed by Dunn’s post hoc test were used to compare female blood meal mass, unfed body mass, fed body mass, and the ratio of blood meal mass to unfed body mass among treatments. Each *P* value is adjusted to account for multiple comparisons. Family-wise significance and confidence level to 0.05. ‘Unmated-Reference’ females were not included in the analysis. We used Fisher’s exact test (two-tailed) to compare olfactometer response rates of unmated and mated females on the same starvation day. Because control filter papers in both sides of the olfactometer resulted in low response rates (<15%), we pooled the controls of the four starvation periods of unmated and mated females. The Kaplan-Meier method was employed to create survival curves from the raw data and a log-rank (Mantel-Cox) test was used to compare the survival curves. Statistical analysis was performed in GraphPad prism v.5.0a.

## Author contributions

A.M.S., Z.C.D., R.S., and C.S. designed the study; A.M.S. conducted the experiments, analyzed the data, and prepared the figures; A.M.S., Z.C.D., and C.S. wrote the manuscript. All authors reviewed and approved the final version of the manuscript.

## Disclaimer

Any opinions, findings, and conclusions or recommendations expressed in this material are those of the authors and do not necessarily reflect the position or the policy of the U.S. Government and no official endorsement should be inferred.

---

**Supplementary Video S1. Expression of a female refusal behavior in** *Cimex lectularius*.

Representative video recording of a 30 days starved Mated-Long female (a female was housed with a fertile male until she died) expressing ‘refusal’ behavior to protect the ectospermalege from a harassing male. The female was taken out of the experimental glass vial for a clearer recording.

